# New tools for automated cryo-EM single-particle analysis in RELION-4.0

**DOI:** 10.1101/2021.09.30.462538

**Authors:** Dari Kimanius, Liyi Dong, Grigory Sharov, Takanori Nakane, Sjors H.W. Scheres

## Abstract

We describe new tools for the processing of electron cryo-microscopy (cryo-EM) images in the fourth major release of the RELION software. In particular, we introduce VDAM, a Variable-metric gradient Descent algorithm with Adaptive Moments estimation, for image refinement; a convolutional neural network for unsupervised selection of 2D classes; and a flexible framework for the design and execution of multiple jobs in pre-defined workflows. In addition, we present a stand-alone utility called MDCatch that links the execution of jobs within this framework with metadata gathering during microscope data acquisition. The new tools are aimed at providing fast and robust procedures for unsupervised cryo-EM structure determination, with potential applications for on-the-fly processing and the development of flexible, high-throughput structure determination pipelines. We illustrate their potential on twelve publicly available cryo-EM data sets.

## 2 Introduction

Structure determination of biological macromolecules through single-particle analysis of cryo-EM images has recently reached a milestone by obtaining atomic resolution reconstructions [1, 2]. With increasing resolutions, the applicability of cryo-EM structure determination continues to improve, and with many inexperienced scientists entering the field, the need for robust, easy to use image processing procedures is increasing. Moreover, atomic resolution structure determination opens up new avenues for cryo-EM structure-based drug design, which often requires high-throughput and automation to enable the screening of many candidate molecules.

The development of user-friendly cryo-EM image processing software has come a long way. Early software packages capable of performing cryo-EM structure determination by single-particle analysis, including SPIDER [3], IMAGIC [4] and the suite of MRC image processing programs [5], were mostly command-line driven and typically relied on extensive user experience to obtain good results. The development of graphical user interfaces (GUIs) and more integrated work flows in the EMAN software [6] reduced this requirement, making cryo-EM image processing accessible to more scientists. Developments in SPARX [7], BSOFT [8], FREALIGN [9] and XMIPP [10] also contributed to improved accessibility. The cryo-EM resolution revolution [11] further accelerated the focus on user-oriented software developments, with new software packages like SPHIRE [12], cisTEM [13] and the commercial cryoSPARC [14] implementing robust and easy-to-use pipelines for cryo-EM structure determination. In addition, overarching software developments like Appion [15] and Scipion [16] facilitated the combination of the different available software packages.

The first release of the RELION software coincided with the appearance of the first prototypes of direct electron detectors that would spark the resolution revolution [17]. RELION introduced a novel empirical Bayesian approach to single-particle analysis, with an explicitly regularised likelihood optimisation target [18]. In the Bayesian framework, parameters for optimal filtering of the reconstruction are inferred from the data, thus removing the need for user expertise to tune related parameters in alternative softwares. Not only did the Bayesian approach lead to higher quality reconstructions; it also represented a step-change in software accessibility that expedited a rapid expansion of the field once direct detectors became commonly available [19].

More recently, automation of large parts of the cryo-EM structure determination pipeline has received increased attention. In particular, various unsupervised protocols for the earlier stages of image processing, including motion correction in movies, contrast transfer function (CTF) parameter estimation and particle picking, have been introduced, for example in FOCUS [20], SCIPION [21], WARP [22], tranSPHIRE [23], SPREAD [24] and cryoFLARE [25]. Automated on-the-fly processing of cryo-EM data allows spotting problems in the data while they are being acquired, thus providing opportunities to change data collection and save valuable time on the microscope. In addition, their standardized procedures lower the barriers for novel users and facilitate the development of high-throughput structure determination pipelines.

This paper describes new tools for single-particle analysis in RELION-4.0 that aim to make unsupervised cryo-EM structure determination faster, more robust and easier to automate.

## 3 Methods

### 3.1 VDAM: Variable-metric Gradient Descent with Adaptive Moments estimation

#### 3.1.1 Regularised likelihood optimization

We briefly recapitulate the regularised likelihood optimization algorithm that underlies classification and refinement procedures in RELION [17, 18].

Let 𝒳 = *x*_1_, …, *x*_*N*_ ∈ ℂ^*L*^ denote the Fourier transforms of the experimental particle images. Each particle image is a noisy 2D projection of a rotated and translated volume, out of an ensemble of unknown volumes with 3D Fourier transforms 𝒱 = *v*_1_, …, *v*_*K*_ ∈ ℂ^*M*^, typically referred to as classes. We assume

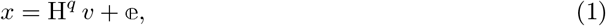

where H^*q*^ ∈ ℂ^*L×M*^ is a complex valued matrix that takes a 2D slice out of the 3D Fourier transform after applying the relevant composite transformation *q* ∈ *Q* := *SE*(3), consisting of a rotation and translation, as well as a (given) contrast transfer function (CTF). We assume uncorrelated Gaussian noise, or 𝕖_*i*_ ∈ ℂ^*L*^ ∼ 𝒞𝒩 (0, *σ*), as well as uncorrelated Gaussian signal, or *v* ∈ ℂ^*M*^ ∼ 𝒞𝒩(0, *τ*), where both 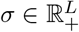 and 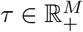 are diagonal co-variance matrices.

We then seek the *maximum a posteriori* (MAP) estimate of 𝒱 by maximizing the following regularised likelihood function:

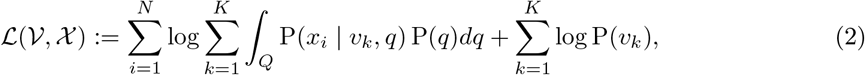

where we have assumed that 𝒳 contains independent observations, and we have marginalised over the nuisance variables, through an integration over *Q* and a summation over *k*. The likelihood term is calculated as 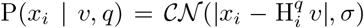; and the prior term as P(*v*) = 𝒞𝒩(0, *τ*). P(*q*) expresses information about the prior probability of the transformations, e.g. a 2D Gaussian distribution for the translations and typically a uniform distribution for rotations.

To find the MAP estimate of 𝒱, we use the Expectation-Maximization algorithm, where we denote each iteration with the index (*n*). Starting from an initial guess, 𝒱^(0)^, we apply a fixed-point iteration approach by fixing 𝒱 and solving ∇_*v*_ ℒ (𝒱, 𝒳) = 0 for the parameters of a particular *v*. First, in the Expectation step we calculate the gradient of (2) with respect to *v*_*k*_:

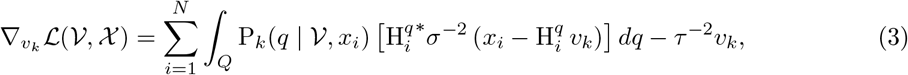

with

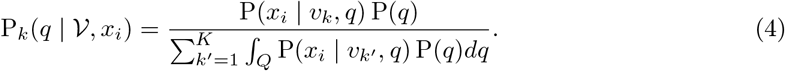

Then, in the Maximization step we solve for the parameters of a particular *v*_*k*_, which yields the closed-form solution:

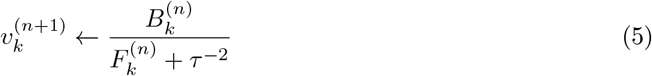

where the division is to be evaluated element-wise, and where

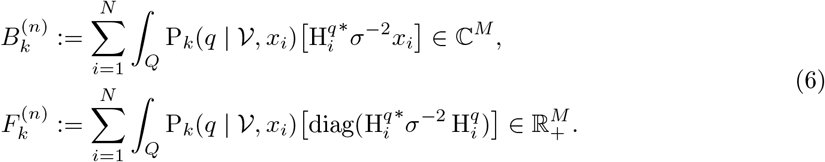

#### 3.1.2 Variable metric gradient descent

Optimisation by gradient descent is an alternative to the Expectation-Maximization algorithm, where the update formula is generally a step in the direction of the negative gradient weighted with the learning rate, *η* ∈ ℝ:

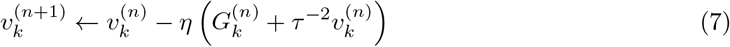

with 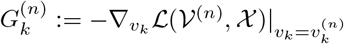, as in (3). In this approach, the summation over *i* in (3) and (6) are typically carried out over a random subset of the data set, called a mini-batch, which is changed at each iteration.

We notice that the Expectation-Maximization algorithm can be viewed as a variable metric gradient descent (VMGD) algorithm, where the gradient in the update formula has been modified with a positive definite projection matrix, *D*^(*n*)^, which changes every iteration [26]. Applying this to the GD update formula in (7) gives

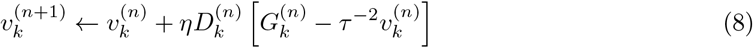

We seek a positive definite matrix *D*^(*n*)^ that equates the gradient descent update to the Expectation-Maximization update. From (5) and (8) we get

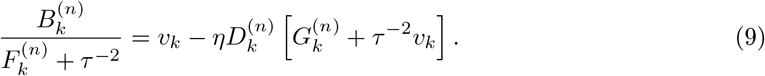

Solving for *D*, yields

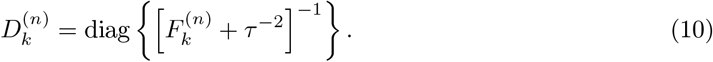

Inserting this into (8) yields the following update formula for the VMGD algorithm:

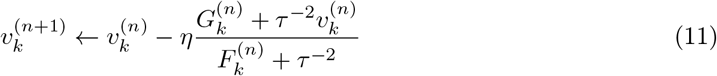

Note that, if the gradient was calculated over the entire data set, *η* can now be set to 1.0, since the gradient is re-scaled by *D* to fit the Expectation-Maximization step size. However, if updates are performed with mini-batches, *η* should still be smaller than 1.0, because the estimated *G* from a subset is noisier. In our implementation, the default values for *η* range 0.1 − 0.9, depending on the stage of reconstruction.

If a Fourier shell correlation (FSC) can be calculated that assesses the signal-to-noise for 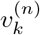, then, using Eqs (9-10) in [17], (11) can be rewritten as:

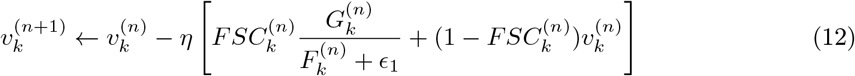

where *ϵ*_1_ is a constant added to improve numerical stability.

In our implementation, we calculate 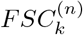 using two separately reconstructed versions of *v*_*k*_, each from one half of the data set 𝒳. Additionally, we define the rescaled gradient *Ĝ* := *G*/(*F* + *ϵ*_1_), which is invariant to the mini-batch size, for small values of *ϵ*_1_.

#### 3.1.3 Adaptive learning rate

We also implemented an adaptive learning rate method, similar to methods commonly used in deep-learning, including Adam [27], Adagrad [28] and ADADELTA [29]. Typically, these methods accumulate two gradient moments, through running averages: *m* ∈ ℂ^*M*^ tracks the momentum of the gradient (first moment), while *u* ∈ ℝ^*M*^ tracks an estimate of the noise or error amplitude in the gradient (second moment). In particular, the Adam optimiser, tracks |*G*|^2^ as the second moment. Instead, we calculate two gradients, 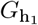 and 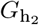, for two separate halves of the data set, h_1_ and h_2_, and accumulate these into two separate first moments. The second moment, which is shared among the two halves, tracks 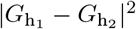. Additionally, we avoid having to track *F* separately by tracking the rescaled gradient *Ĝ* instead. Thereby, we consider the following three running averages for each *k*:

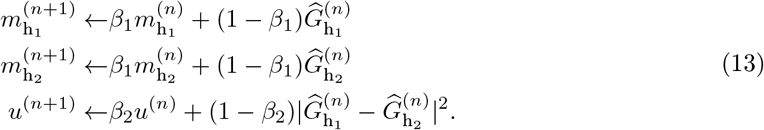

The update formula for each half data set then becomes:

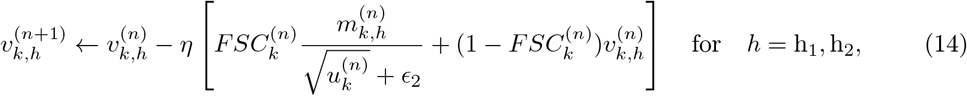

where *ϵ*_2_ is a suitably small constant. In our implementation, *β*_1_ = 0.9 and *β*_2_ = 0.999.

Excluding the regularisation term in (12) from the running averages in (13) serves to decouple the effects of regularisation and the learning rate, which has been shown to improve convergence efficiency for the Adam optimiser [30].

#### 3.1.4 Replacing inactive classes

In previous releases of RELION, especially in 2D classification, many classes would converge to contain no or very few particles. This represented a waste of computational resources and often resulted in suboptimal classification of structural variability in the data. To address this issue we here propose an algorithm that, throughout the gradient optimisation, substitutes classes with too small likelihood probabilities with classes that exhibit large variability. This approach is inspired by methods used in the class of artificial neural network algorithms known as self-organizing maps and neural gases [31]. At the end of each iteration, before the gradient update is applied, we select class *a*, which is the class with the smallest likelihood probability, P(𝒳 | *v*_*a*_). Next, we select class *b*, which is the class with the largest 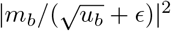. The following modified update formula is then used for the two classes:

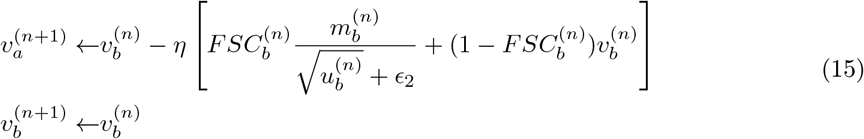

Similarly, the moments of class *a* are substituted with that of class *b*. Substitutions are only performed if P(𝒳 | *v*_*a*_) < *ρ/K*, where *ρ* is a user-defined constant, called the *class inactivity threshold* (see section 4.3).

#### 3.1.5 Implementation details

In previous releases of RELION, the initial references for 2D classification and 3D initial model generation were initialised from averages of random subsets of the particles in random orientations. In RELION-4.0, the default is to start from randomly positioned Gaussian blobs that are different for each class. The primary purpose of this initialisation is to diversify the classes early in the classification, thus leading to faster convergence and the recovery of more class variability.

To further reduce computational costs for both 2D classification and 3D initial model generation, VDAM optimisations are started with a high learning rate of 0.9. By default, the learning rate is then gradually reduced to 0.3 for 2D classification and to 0.5 for 3D initial model generation. In addition, calculations are started from relatively small mini-batches: 0.5% of the total data set size (with a minimum of 200 particles, and a maximum of 10,000 particles for 2D classification and a maximum of 5,000 for 3D initial model generation). After the initial stage the learning rate is reduced to 0.3 for 2D classification and to 0.5 for 3D initial model generation, and the mini-batch size is increased to 5% and 10% of the data set size, respectively (with a minimum of 1,000 particles, and a maximum of 100,000 particles for 2D classification and 50,000 particles for 3D initial model generation).

Although most of the data sets tested in this paper converge after 100 mini-batches, in order to obtain good results for a larger number of data sets, we set the default on the GUI to 200 mini-batches, and ran all calculations in this paper using 200 mini-batches. In this way, the total number of passes through the entire data set, i.e. epochs, is five or less, resulting in a major speed-up compared to the 25 full iterations that are done by default using the EM algorithm. In the final iteration, a final pass through the entire data set is performed, where only P(𝒳 | *k*) for each class *k* is calculated, further saving time compared to a normal epoch. In addition, we noticed that the VDAM algorithm is less sensitive to truncation of the marginalisation (i.e. skipping those orientations from the integral in eq. (2) with low probabilities) than the EM algorithm, leading to additional increases in speed, in particular during the early stages of refinement.

### 3.2 Class ranker: automated 2D class selection

The selection of particles that give rise to 2D class average images with recognisable protein features is often used to discard suboptimal particles from cryo-EM data sets. The selection of suitable 2D classes was done interactively in previous releases of RELION. RELION-4.0 contains a new program called `relion_class_ranker` that automates 2D class selection. This program predicts a score for each class by combining the output of a convolutional neural network that acts on the 2D class average images with 18 features (Figure 1A,B).

**Figure 1:**
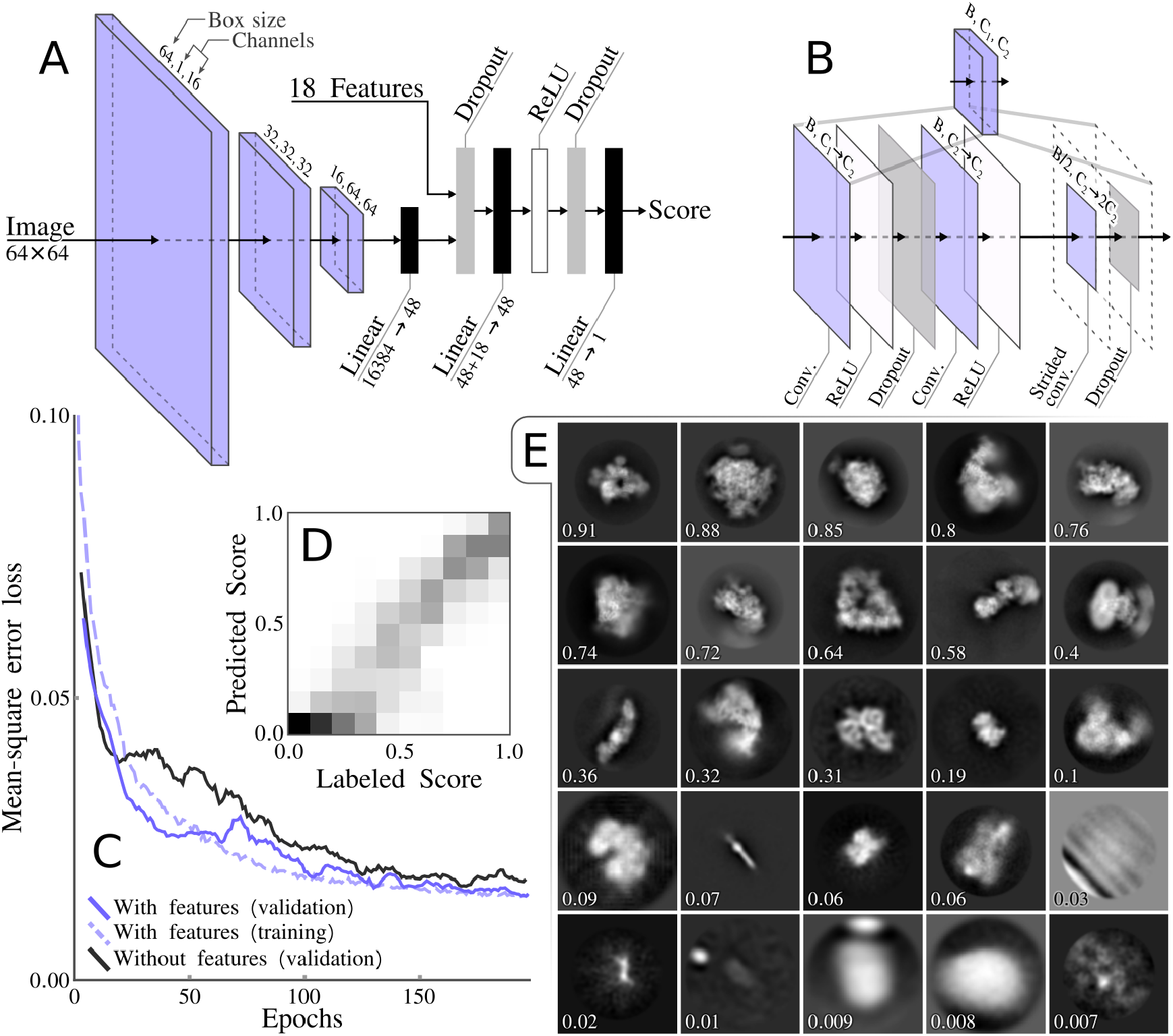
Class ranker neural network architecture and results. A) The overall architecture of the scoring network, which consists of three CNN blocks and a final feed forward network that also incorporates the 18 features. B) The CNN block architecture that incorporates three convolutional layers. The initial convolutional layer, maps the input channels count *C*_1_ to the intermediate channels count *C*_2_ and the final layer preforms a down sampling of the box size through a strided convolution and doubles the number of channels. C) The mean-square error loss during training, comparing with and without features. D) A confusion matrix showing labeled scores versus predicted scores, with bins of 0.1. E) Example of classes with their predicted score.

The convolutional neural network takes as input individual 2D class average images, cropped to contain only the area defined by the circular mask used in the 2D classification, and re-scaled to 64 × 64 pixels. The feature vector is calculated for each class from RELION’s metadata of the 2D classification job, including the estimated accuracies of rotational and translational alignments, the estimated resolution (1*/d* in 1*/*Å) and a so-called weighted resolution, which is calculated as *d*^2^*/*ln*N*, where *N* is the number of particles assigned to the class. It also contains features that are calculated from the 2D class average images, in particular the first to fourth moments of density values inside an automatically determined mask for the protein region, the solvent region, and for a ring around the outer diameter of the mask that has been applied to the 2D class average images. The combined output from the convolutional neural network and the feature vector is passed through two fully-connected layers, with non-linear (ReLU) activation functions between the layers, to predict a single, floating point value, score for each 2D class.

The network in the `relion_class_ranker` program was trained on 18,051 2D class average images from 233 RELION 2D classification jobs that were performed at the MRC-LMB over a period of approximately four years. Each of the jobs was assigned a job score, ranging from zero to one, and within jobs the class averages images were manually divided into four categories depending on their quality. For each 2D class, the combination of its job score, its category assigned, and its estimated resolution compared to the best resolution in its 2D classification job, were used to calculate a target class score, ranging from zero to one. The target scores were intended to represent a ranking over all classes in the training set, with a score of one representing the best classes from the best 2D classification jobs, and a score of zero representing the worst classes. The network was implemented and optimised with the Adam optimiser [27] for 200 epochs in pytorch [32], using a mean-squared error between predicted and assigned scores. All 18,051 class average images, plus their metadata from the 2D classification jobs and their assigned class scores are publicly available through the EMPIAR data base (entry-ID 10812). The code used to optimise and execute the neural network are available from the RELION github pages.

### 3.3 Schemes: planning and execution of multiple jobs

RELION’s pipeliner organizes the execution of RELION jobs, which represent individual tasks, and often the execution of an individual command-line program, in the overall structure determination workflow. The pipeliner also keeps track which jobs’ output files are used as input for other jobs, thus building a directional graph of the processing workflow [33].

RELION-4.0 implements new functionality to plan the execution of multiple jobs in advance, including functionality to execute branched decision trees, where decisions to follow one branch of sequential jobs or another are made on-the-fly. A series of planned jobs, possibly including multiple branches, is called a *Scheme*.

To allow for flexibility in the design of the execution of multiple jobs, Schemes implement different types of variables: *stringVariables, booleanVariables* and *floatVariables*. The values of these variables can be changed through the execution of so-called *Operators* that form part of the Scheme framework. Multiple Operators have been implemented, for example to perform simple mathematical operations on floatVariables; to perform logical operations on booleanVariables; and to perform text modifications on StringVariables. In addition, Operators exist for reading metadata values from RELION star files; for file handling operations; for sending emails and for waiting a pre-determined amount of time. A full description of available Operators is available from the RELION documentation, which has been rewritten, and is available from: https://relion.readthedocs.io/en/release-4.0/.

Schemes can be thought of in terms of a directional graph, where the nodes of the graph are either jobs or Operators. *Edges* connect two subsequent nodes, while *Forks* connect one input node to two possible output nodes. Each Fork has an associated booleanVariable, whose value determines which of the two output nodes is chosen upon execution of the Scheme. The topology of the graph inside a Scheme can be cyclical, thereby enabling repetitive execution of jobs inside loops.

Schemes are defined by a `scheme.star` file that describes the different jobs, Operators, Variables, Edges and Forks. The `scheme.star` file is stored inside a `Schemes/schemename` directory, which itself is inside the standard RELION Project directory. The Scheme directory also contains subdirectories for each of the RELION jobs that form part of the Scheme. These subdirectories each contain a file called `job.star` that contains the parameter values for that job. The Scheme framework is flexible, in that users can define their own Schemes by manually editing the corresponding star files. The `job.star` files for individual jobs can also be saved through the Jobs menu on the RELION-4.0 main GUI.

Schemes are executed through the `relion_schemer` program, which launches the jobs, and keeps track of the current status of the Scheme and the values of all its variables. It can also be used to abort a running Scheme, to change its current variables or the parameters of its RELION jobs, and to re-start from the point where it was previously aborted. If any RELION job parameters were changed, the `relion_schemer` program will re-execute those jobs from scratch, whereas jobs that are unaffected by the changes will continue from where they were halted.

RELION-4.0 includes two example Schemes, called `prep` and `proc`. The `prep` Scheme imports micrograph movies and performs motion correction and CTF estimation. The `proc` Scheme selects micrographs based on their estimated CTF resolution limit, performs automated particle picking (using either RELION’s Laplacian-of-Gaussian (LoG) approach or Topaz [34]), 2D classification, automated 2D class selection, 3D initial model generation and 3D refinement. A flowchart of both Schemes, depicting all corresponding RELION jobs and Scheme Operators is shown in Figure 2.

**Figure 2:**
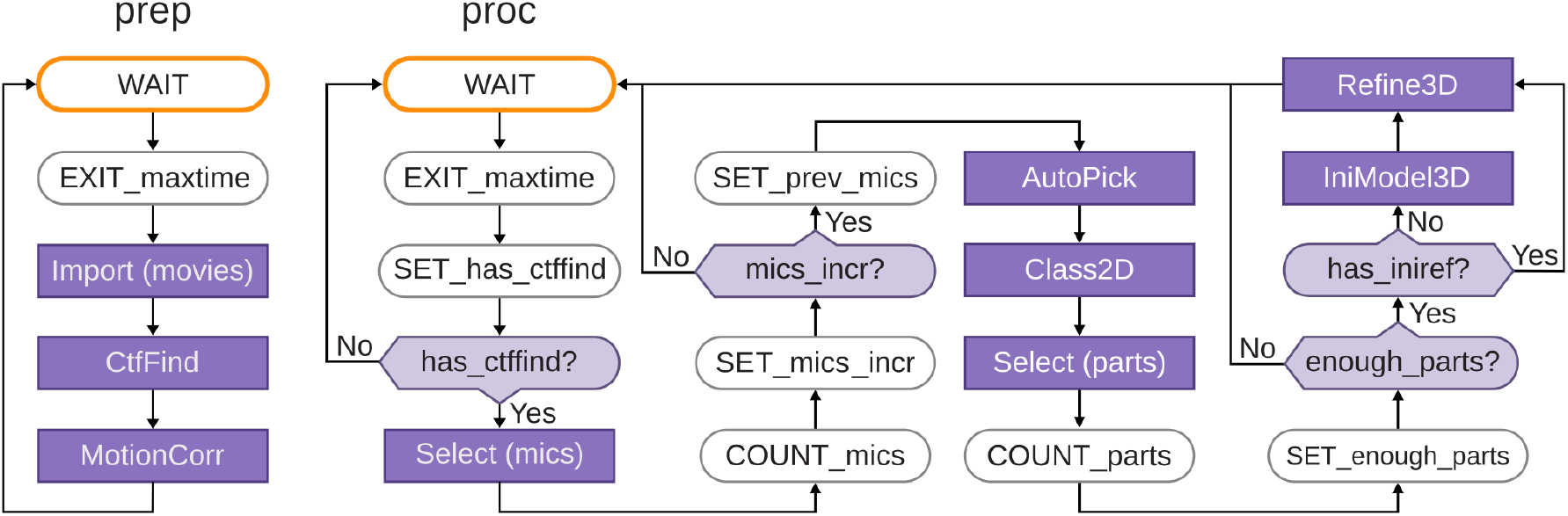
Schematics of the `prep` and `proc` Schemes that form part of the `relion_it.py` approach for automated, on-the-fly processing. Scheme operators are shown with rounded boxes, RELION jobs with grey boxes; edges with arrows and forks with diamond-shapes. For forks, the BooleanVariable that controls its outcome is indicated in the centre of the diamond. The `WAIT` operator waits for a defined time since it was last executed; the `EXIT_maxtime` operator terminates the Scheme after a defined time since the Scheme was started; the `SET_has_ctffind` operator sets BooleanVariable `has_ctffind` to true if the STAR file generated by the CtfFind job of the `prep` Scheme exists; the `COUNT_mics` operator sets the current number of micrographs to the number selected in the job above it; the `SET_mics_incr` sets BooleanVariable `mics_incr` to true if the current number of selected micrographs is larger than the previous number of micrographs (which is initialised to zero); the `SET_prev_mics` operator sets the previous number of micrographs to the current number of selected micrographs; the `COUNT_parts` operator sets the current number of particles to the number of selected particles in the job above it. The `SET_enough_parts` operator sets BooleanVariable `enough_parts` to true if the current number of selected particles is larger than a user-specified minimum.

The python script `relion_it.py`, which already existed in RELION-3, has been modified to work with Schemes in RELION-4.0. The modified script launches a GUI to gather parameter input from the user and then executes the `prep` and the `proc` Schemes to process cryo-EM data sets in an unsupervised manner. In addition, a new GUI called `relion_schemegui.py` has been written to facilitate the monitoring of Schemes during their execution, as well as their aborting, changing of variables, and re-starting.

### 3.4 MDCatch: integration with the microscope

To simplify launching of on-the-fly image processing and minimize user input errors we implemented MDCatch, a graphical tool that extends `relion_it.py` functionality by linking microscope data acquisition with the execution of RELION-4.0 Schemes. MDCatch is written in Python3 and PyQt5 and provides a simple GUI (Figure 3) that fetches acquisition metadata from a running EPU (Thermo Fisher Scientific) or SerialEM [35] session, and launches a predefined image processing pipeline. Besides RELION-4.0 Schemes, MDCatch also works with Scipion 3 workflows [21]. MDCatch supports parsing of metadata from different file formats (EPU’s XML, SerialEM’s MDOC, MRC, TIF) and associates this information with other microscope parameters (detector type, MTF, gain reference etc.) that can be configured in advance. In cases where existing metadata is not sufficient, users can manually input missing information. MDCatch was designed to be installed on a computer that has access to both the raw data (movies) and its associated metadata, e.g. an EPU session folder. For SerialEM data acquisitions, both movies and MDOC metadata files are expected to be in the same directory.

**Figure 3:**
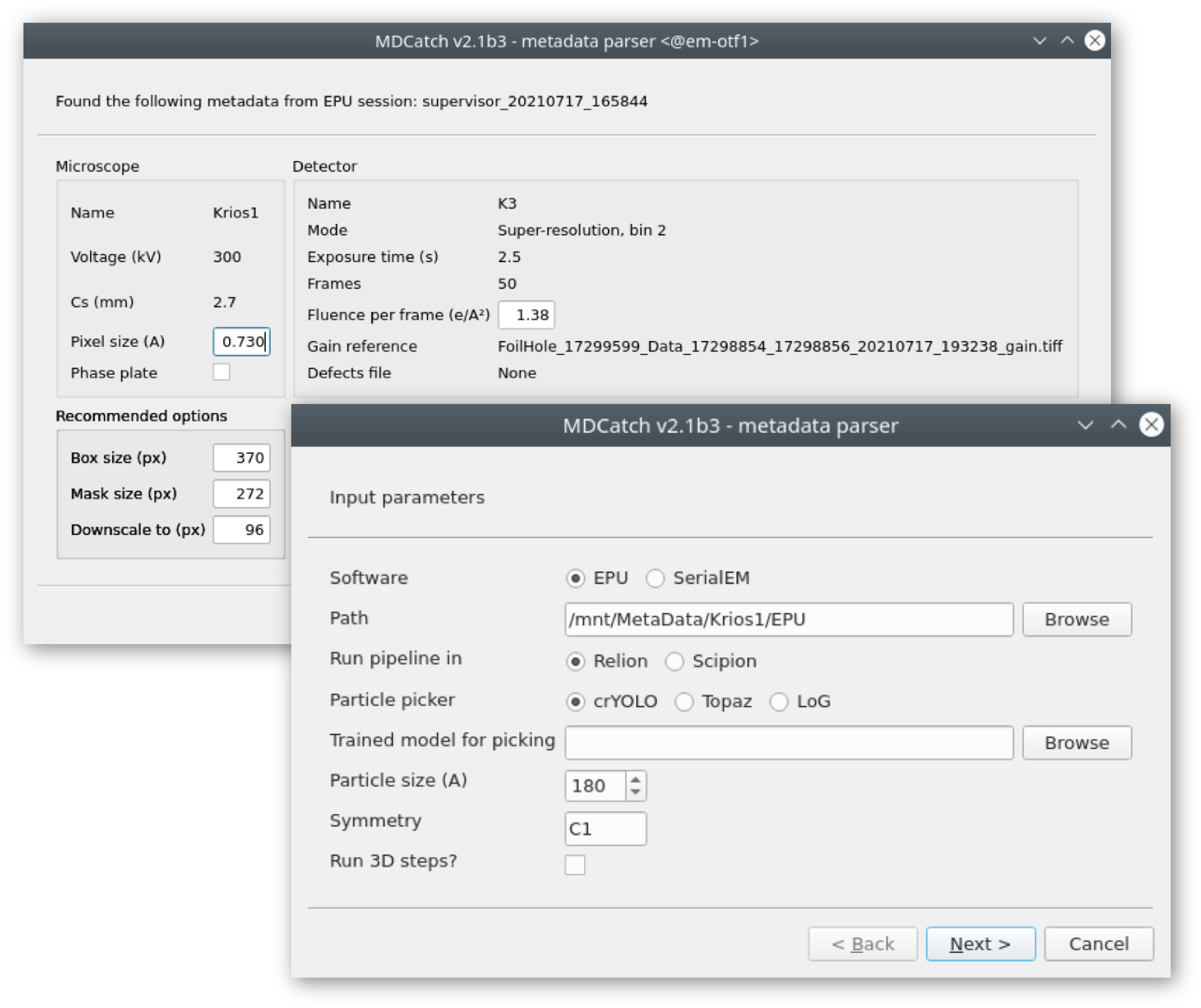
GUI of the MDCatch utility for automated fetching of microscope metadata and launching of on-the-fly image processing.

Together with MDCatch we provide example pipeline templates for both RELION and Scipion, including the `prep` and `proc` Schemes described above. By default, users are presented with a choice between three particle pickers: the Laplacian-of-Gaussian (LoG) approach in RELION, crYOLO [36] or Topaz [37]. Upon execution of MDCatch, the fetched metadata are saved in a text file and the image processing progress can be monitored with existing project utilities in RELION or Scipion.

MDCatch is distributed separately from RELION-4.0, under a free, GPLv3 software license, and its code and documentation are available at https://github.com/azazellochg/MDCatch.

## 4 Results

### 4.1 Optimisation of the neural network in relion_class_ranker

During exploration of network architecture and optimisation procedures, 5,850 2D class average images, from 73 2D classification jobs, were set aside as a validation set to monitor overfitting.

Because the final network architecture and optimisation procedure did not induce noticeable amounts of overfitting (Fig 1C), a final optimisation round was performed using all 18,051 classes. The resulting network had a mean-square error loss of 0.015 on the predicted class scores. Optimisation of a network where all feature values were set to zero led to a mean-square error loss of 0.017, indicating that the features provided useful information in the scoring process. Analysis of 2D histograms of the assigned and predicted class scores (Fig 1D) and manual assessment of the predicted scores (Fig 1E) confirmed that the final network produces useful predictions over the full range of assigned class scores. The optimised network was further tested as part of the automated processing of twelve test data sets through the Schemes and `relion_it.py` approach, as described below.

### 4.2 Automated processing with Schemes and relion_it.py

To test the procedures for automated single-particle analysis in RELION-4.0, we processed twelve data sets from the EMPIAR data base [38], using default parameters from `relion_it.py`, except for the experiment-specific parameters and the particle diameter as shown in Table 1. The twelve data sets were selected at the start of the project; no data sets were added or removed during the project. Motion correction for movies of these data sets were performed in RELION’s own implementation of the MotionCor2 algorithm [39]. CTF estimation was performed in CTFFIND4 [40]. Motion-corrected micrographs and extracted particles were saved in IEEE 754 16-bit float MRC format (mode 12), a new feature in RELION-4.0 to save a factor of two in required disk space. The `proc` Scheme was used for automatically processing the data, up to 3D initial model generation and refinement of down-scaled particle images.

**Table 1:**
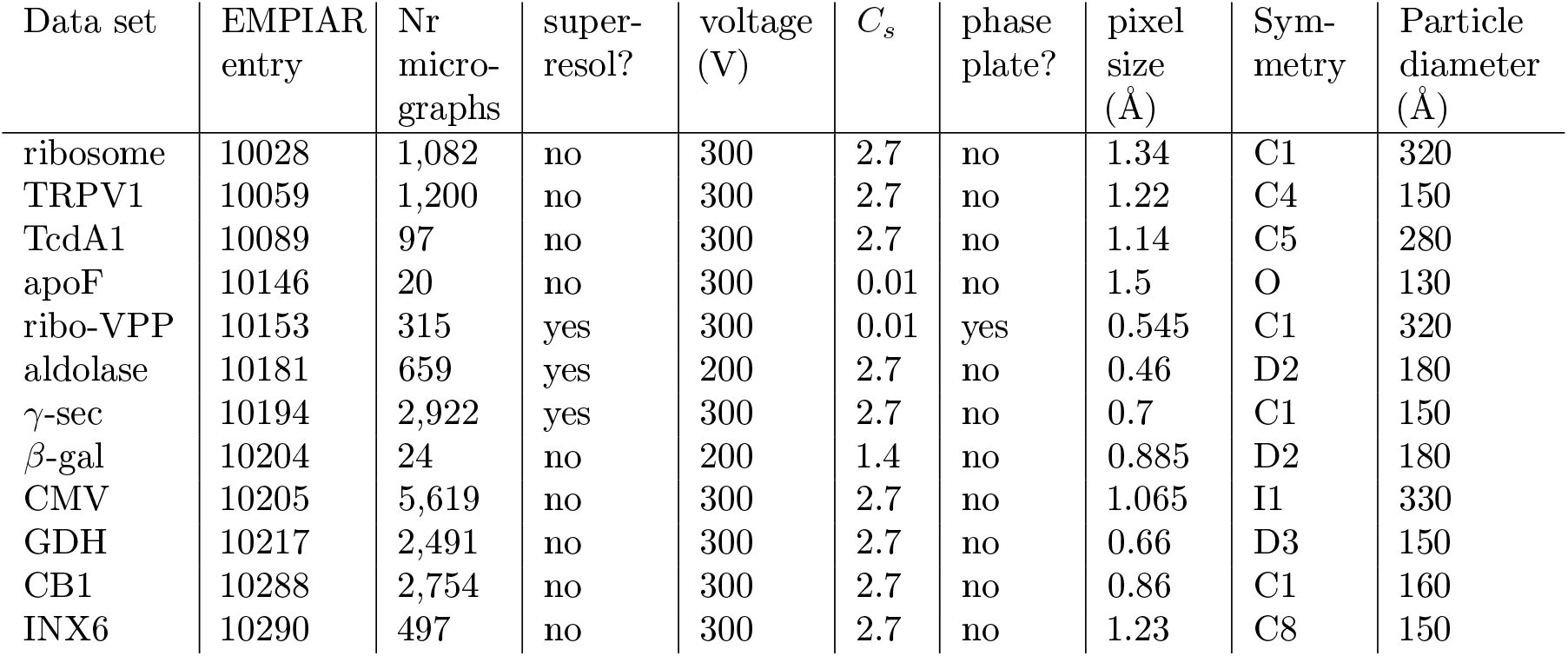
Test data set characteristics. ribosome: Plasmodium falciparum 80S ribosome [41] TRPV1: transient receptor potential channel V1 [42]; TcdA1: Tripartite Tc toxin subunit A [12]; apoF: apoferritin [43]; ribo-VPP: ribosome collected on a Volta phase plate [44]; aldolase: Rabbit muscle aldolase [45] *γ*-sec: *γ*-secretase [46]; *β*-gal: *β*-galactosidase [47]; CMV: cowpea mosaic virus [48]; GDH: glutamate dehydrogenase [49]; CB1: cannabinoid receptor 1G [50]; INX6: innexin-6 hemichannel [51];

| Data set | EMPIAR entry | Nr micro-graphs | super-resol? | voltage (V) | $C_s$ | phase plate? | pixel size (Å) | Symmetry | Particle diameter (Å) |
| --- | --- | --- | --- | --- | --- | --- | --- | --- | --- |
| ribosome | 10028 | 1,082 | no | 300 | 2.7 | no | 1.34 | C1 | 320 |
| TRPV1 | 10059 | 1,200 | no | 300 | 2.7 | no | 1.22 | C4 | 150 |
| TcdA1 | 10089 | 97 | no | 300 | 2.7 | no | 1.14 | C5 | 280 |
| apoF | 10146 | 20 | no | 300 | 0.01 | no | 1.5 | O | 130 |
| ribo-VPP | 10153 | 315 | yes | 300 | 0.01 | yes | 0.545 | C1 | 320 |
| aldolase | 10181 | 659 | yes | 200 | 2.7 | no | 0.46 | D2 | 180 |
| γ-sec | 10194 | 2,922 | yes | 300 | 2.7 | no | 0.7 | C1 | 150 |
| β-gal | 10204 | 24 | no | 200 | 1.4 | no | 0.885 | D2 | 180 |
| CMV | 10205 | 5,619 | no | 300 | 2.7 | no | 1.065 | I1 | 330 |
| GDH | 10217 | 2,491 | no | 300 | 2.7 | no | 0.66 | D3 | 150 |
| CB1 | 10288 | 2,754 | no | 300 | 2.7 | no | 0.86 | C1 | 160 |
| INX6 | 10290 | 497 | no | 300 | 2.7 | no | 1.23 | C8 | 150 |

Table 2 gives an overview of the results. Only micrographs with resolutions beyond 6 Å, as estimated by CTFFIND-4, were included in the processing. For all data sets, except for the ribosome data set collected with a phase plate (ribo-VPP; EMPIAR-10153), particle picking using the pretrained model in Topaz yielded reasonable results. For the ribo-VPP data set, the unusually strong contrast in the phase plate images yielded suboptimal results in Topaz, and we used the LoG-picker in RELION instead. All particles were extracted in the box sizes suggested by `relion_it.py`, i.e. 1.5 times the particle diameter, and downscaled to pixel sizes in the range of 2.8-3.5Å (with the exact pixel size depending on favourable downscaled image sizes for the fast Fourier Transform algorithm). The extracted particles were subjected to 2D classification with 100 classes, using the VDAM algorithm, followed by automated class selection in `relion_class_ranker` with a default minimum class score of 0.15. Finally, the selected particles were subjected to 3D initial model generation in symmetry group C1, again using the VDAM algorithm, with three classes, followed by 3D auto-refinement of the largest class after automated detection and alignment of the symmetry axes.

**Table 2:** Summary of automated processing of the test data sets. Data sets are as defined in Table 1. LoG: Laplacian of Gaussian; * For the TRPV1 and aldolase data sets, initial model generation often get stuck in a local minima, leading to an incorrect map. This issue is observed occasionally also for GDH. See section 4.5 for more details.

| data set | Selected micrographs | Picking job | Picked particles | Selected particles | Correct map? |
| --- | --- | --- | --- | --- | --- |
| ribosome | 1,081 | Topaz | 188,844 | 161,194 | yes |
| TRPV1 | 1,199 | Topaz | 520,020 | 184,836 | yes* |
| TcdA1 | 97 | Topaz | 19,479 | 11,818 | yes |
| apoF | 19 | Topaz | 2,282 | 0 | no |
| ribo-VPP | 298 | LoG | 22,895 | 17,606 | yes |
| aldolase | 659 | Topaz | 369,520 | 186,327 | yes* |
| γ-sec | 2,873 | Topaz | 1,665,273 | 601,160 | yes |
| β-gal | 24 | Topaz | 8,173 | 4,631 | yes |
| CMV | 5,526 | Topaz | 492,981 | 266,621 | yes |
| GDH | 2,427 | Topaz | 476,563 | 400,849 | yes* |
| CB1 | 2,751 | Topaz | 1,196,915 | 470,293 | yes |
| INX6 | 497 | Topaz | 276,868 | 76,446 | yes |

For all data sets, except the apoferritin data set (EMPIAR-10146), 2D classification with the VDAM algorithm (see section 4.3) provided adequate information to assess the quality of the data and the class ranking successfully identified suitable 2D class averages (see section 4.4). Dense packing of the apoferritin particles in the micrographs caused the appearance of density for neighbouring particles in the 2D class averages, which resulted in too low class scores. Because only 294 apoferritin particles were selected, no 3D model generation was attempted. For all other data sets, correct reconstructions could be obtained in a fully automated manner with resolutions close to the Nyquist limit for the downscaled particles (but also see section 4.5).

### 4.3 2D classification with the VDAM algorithm

Figure 4 shows two example comparisons between 2D classifications with the VDAM and the EM algorithms: for the GDH and CB1 data sets. For each run, we manually selected the best classes (highlighted in purple in Figure 4 for GDH and CB1), and subsequently used the corresponding sets of particles for 3D auto-refinement to asses the relative quality of the classified subsets. Computations were performed on an Intel Xeon Gold 6242 and four NVIDIA GeForce RTX 2080Ti GPUs. All VDAM calculations were performed with the default class inactivity threshold of 0.1.

**Figure 4:**
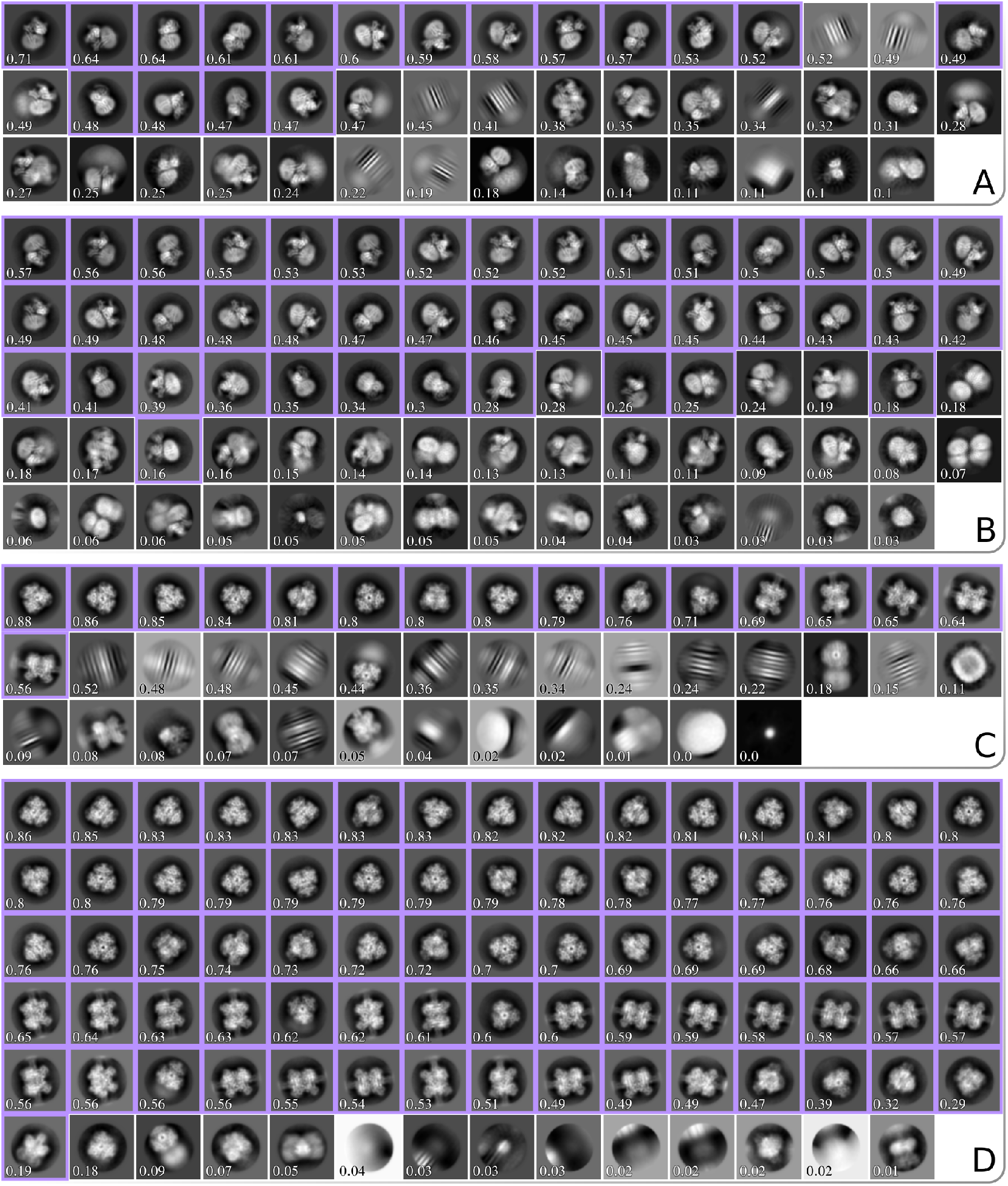
All significant 2D class averages from four different classification runs. Panel A and B show results for the GDH data set classified using the EM and VDAM algorithm, respectively. Panel C and D show results for the CB1 data set classified using the EM and VDAM algorithm, respectively. Classes are sorted according to their score from the `relion_class_ranker program,` which is also shown for each class. Classes that were manually selected for subsequent 3D auto-refinement are highlighted in purple.

Table 3 shows the results for five test data sets. Each run consists of 25 EM and 200 VDAM iterations, which corresponds to 25 and approximately 6 epochs, respectively. The final epoch for the VDAM algorithm is only performed to assess particle class assignment, and is thus faster. On average, the VDAM algorithm is a factor of 5 faster compared to the EM algorithm for 2D classification, without affecting the quality of the selected particles, as measured by the final resolution after auto-refinement.

**Table 3:**
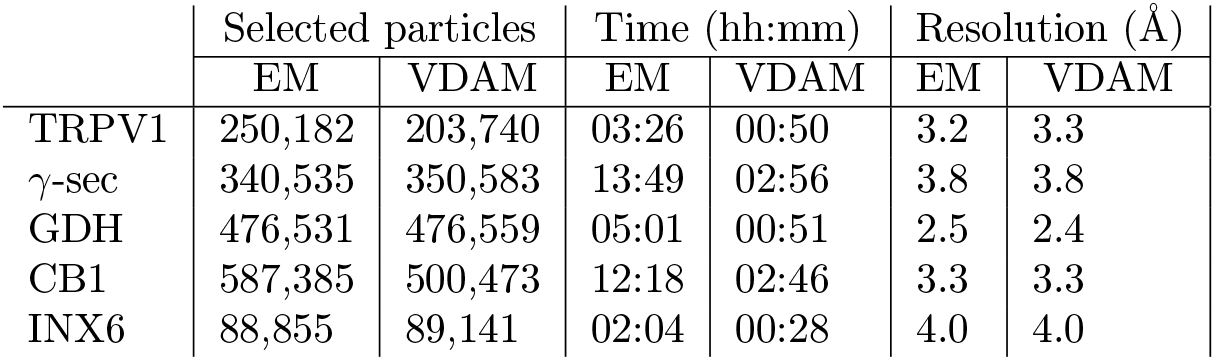
Comparison between the two algorithms for 2D classification. Good classes were selected manually, and the selected subset was further processed in auto-refine. The number of manually selected particles are shown for each data set and algorithm, as well as the execution time for 2D classification and the final resolution achieved with auto-refine using the subset from the respective subsets.

|  | Selected particles |  | Time (hh:mm) |  | Resolution (Å) |  |
| --- | --- | --- | --- | --- | --- | --- |
|  | EM | VDAM | EM | VDAM | EM | VDAM |
| TRPV1 | 250,182 | 203,740 | 03:26 | 00:50 | 3.2 | 3.3 |
| γ-sec | 340,535 | 350,583 | 13:49 | 02:56 | 3.8 | 3.8 |
| GDH | 476,531 | 476,559 | 05:01 | 00:51 | 2.5 | 2.4 |
| CB1 | 587,385 | 500,473 | 12:18 | 02:46 | 3.3 | 3.3 |
| INX6 | 88,855 | 89,141 | 02:04 | 00:28 | 4.0 | 4.0 |

### 4.4 Automated 2D class selection with relion_class_ranker

The predicted class score of the `relion_class_ranker` program was designed to be on an absolute scale, ranging from a value of zero for useless classes to a value of one for the best classes from the best data sets. Therefore, because some data sets are better than others, the threshold for class selection may need to be adjusted in line with the expected quality of the 2D class average images for a given data set. Table 4 shows an evaluation of the quality of the particle selection for all twelve test data sets, by comparing the selections based on the indicated class score threshold (*t*) with a manual selection of suitable classes. Quality is measured in terms of the false positive rate (FPR) and the false negative rate (FNR) of the particles from the selected 2D classes, where the particles from the manually selected classes are considered the correct ones. To reflect that the threshold value may be changed based on the expected quality of each data set, besides reporting the results for the default score threshold of 0.15 used in `relion_it.py`, this table also shows the results for a freely chosen, i.e. supervised threshold value (*t* = *T*) for each data set.

**Table 4:** Analysis of the automated 2D class selection. The second, third and seventh column show the number of particles in the selected classes after manual selection of the classes, automated class selection with a threshold of 0.15, and automated class selection with a supervised threshold, respectively. The sixth column shows the supervised choice for the threshold (t=T). FPR: false positives rate, i.e. number of particles selected by the class ranking procedure, but not by manual selection, divided by the number of selected particles in the manual selection; FNR: false negative rate, i.e. number of particles selected by manual selection, but not by the class ranking procedure, divided by the number of selected particles by manual selection.

| data set | manual<br>selection | class<br>ranker<br>($t=0.15$) | FPR<br>($t=0.15$) | FNR<br>($t=0.15$) | super-<br>vised<br>$t=T$ | class<br>ranker<br>($t=T$) | FPR<br>($t=T$) | FNR<br>($t=T$) |
| --- | --- | --- | --- | --- | --- | --- | --- | --- |
| ribosome | 142,373 | 161,194 | 0.13 | 0.00 | 0.5 | 139,249 | 0.00 | 0.02 |
| TRPV1 | 268,651 | 184,836 | 0.03 | 0.34 | 0.055 | 312,980 | 0.21 | 0.05 |
| TcdA1 | 12,529 | 11,818 | 0.00 | 0.06 | 0.05 | 124,69 | 0.00 | 0.01 |
| apoF | 1,949 | 124 | 0 | 0.94 | 0.02 | 1,874 | 0.04 | 0.08 |
| ribo-VPP | 17,768 | 17,606 | 0.00 | 0.01 | 0.1 | 17,740 | 0.00 | 0.00 |
| aldolase | 170,641 | 186,327 | 0.15 | 0.05 | 0.129 | 192,282 | 0.15 | 0.02 |
| γ-sec | 416,231 | 601,160 | 0.44 | 0.00 | 0.21 | 414,086 | 0.10 | 0.10 |
| β-gal | 6,165 | 4,631 | 0.00 | 0.25 | 0.035 | 6,140 | 0.03 | 0.03 |
| CMV | 229,943 | 266,621 | 0.16 | 0.00 | 0.32 | 231,885 | 0.01 | 0.00 |
| GDH | 416,197 | 400,849 | 0.00 | 0.04 | 0.08 | 421,980 | 0.01 | 0.00 |
| CB1 | 420,739 | 470,293 | 0.15 | 0.03 | 0.225 | 415,574 | 0.02 | 0.03 |
| INX6 | 113,861 | 76,446 | 0.00 | 0.33 | 0.0498 | 140,050 | 0.24 | 0.01 |

For the ribosome, CMV, CB1 and *γ*-sec data sets, a higher threshold than the default leads to better results, although only for the *γ*-sec data set the FPR is higher than 0.2 using the default threshold. For the TRPV1, apoF, *β*-gal and INX6 data sets, a lower threshold yields better results, with the default threshold yielding FPRs of 0.25 or higher. Nevertheless, as pointed out above, even when using the default threshold of 0.15, the particle selection for all data sets, except apoF, allowed *de novo* reconstruction of a correct 3D map. Using the supervised threshold, for all data sets the FPR and the FNR are below 0.25.

### 4.5 Initial 3D model generation with the VDAM algorithm

To evaluate the overall performance of the VDAM algorithm for 3D initial model generation, we performed ten repeats of 3D initial model jobs for the particle sets that were selected automatically by the `relion_it.py` approach of the five data sets shown in Figure 5. Following the `relion_it.py` approach in RELION-4.0, we used the VDAM algorithm with three classes and selected the most populated class after 200 iterations. For each run, we used the selected model as initial reference for subsequent auto-refinement, and used Fourier shell correlation of the refined structure to the known target structure, after manual alignment in Chimera [52], as a metric to distinguish between successful and unsuccessful runs.

**Figure 5:**
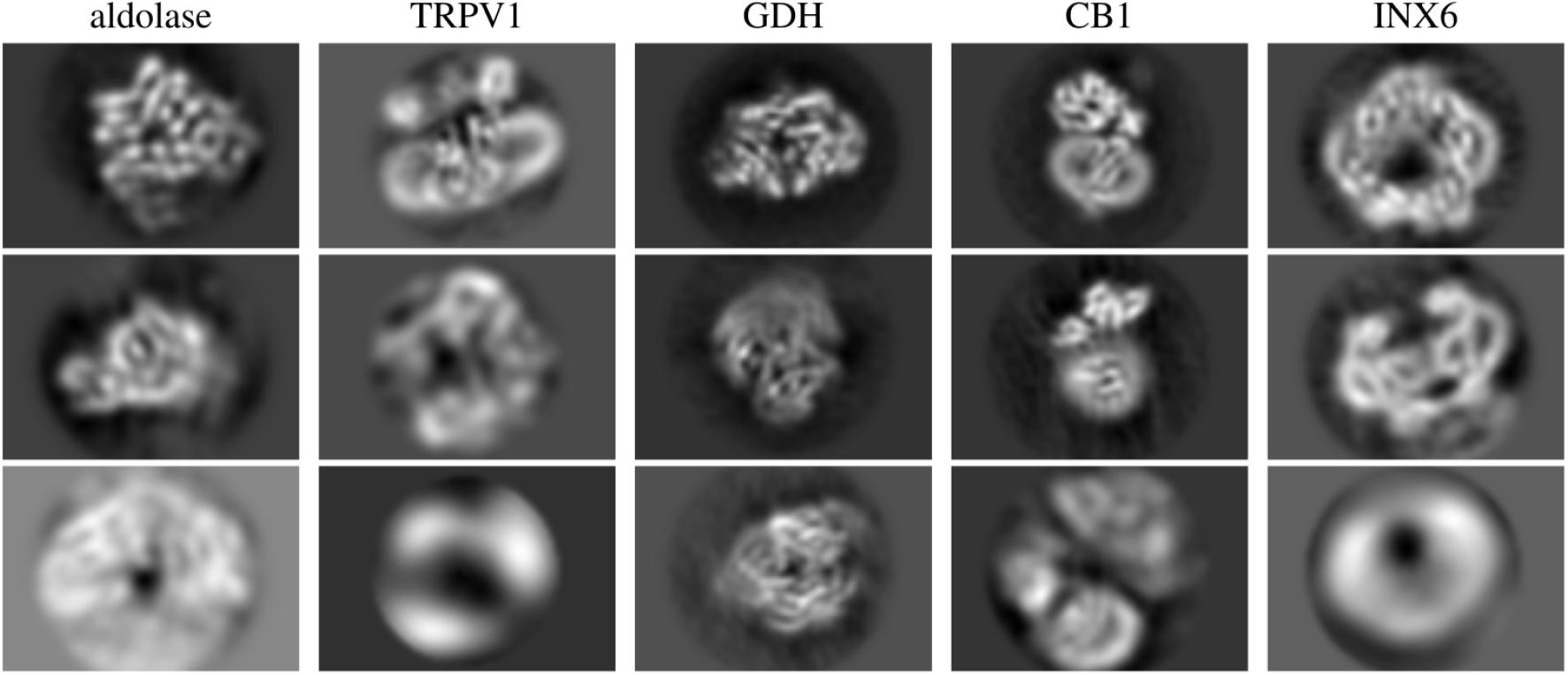
Central slices of initial model reconstruction with 3 classes using VDAM algorithm for 5 data sets.

Figure 5 shows examples of central slices of initial model reconstructions that converged to the target structures for the five data sets. For the GDH, CB1 and INX6 data sets, initial model generation was successful for 8, 9 and 10 out of the 10 runs performed, respectively. For the aldolase data sets, 5 out of the 10 VDAM calculations were successful. Convergence within 200 iterations was primarily hindered by heterogeneity, since a large subset of the data set consisted of similar but incomplete particles. Since the automated procedure picks the most populated class, sometimes the target structure is missed as it comes in at second place. The target structure, exhibiting D2 symmetry, makes up about 50% of the data set after the automated 2D class selection. Manual selection might be a necessity to acquire a good final reconstruction for this data set. For the TRPV1 data set, only 4 out of 10 runs were successful. In this case, angular alignment posed the primary issue, possibly due to the relatively large membrane patch.

## 5 Discussion

In this paper, we present new features for single-particle analysis in RELION-4.0: the VDAM algorithm for image refinement, the `relion_class_ranker` program for automated 2D class selection, and the Schemes framework with an updated `relion_it.py` approach for unsupervised execution of workflows. In addition, we describe the separately distributed MDCatch program that collects metadata from the microscope to facilitate on-the-fly data processing. The RELION-4.0 release also introduces new approaches to sub-tomogram averaging, and a tighter integration of its pipeline approach with the CCP-EM software suite [53]. These two developments will be described elsewhere.

By iterating through the data set fewer times, the VDAM algorithm provides substantial increases in speed compared to the EM algorithm. For example, we illustrate that 2D classification with the VDAM algorithm is up to six times faster than the EM algorithm, with larger gains in speed observed for larger data sets. Even larger speed-ups may be obtained by reducing the default number of mini-batches from 200 to 100. Although performing fewer iterations may affect the quality of the results for difficult data sets, the additional speed-ups obtained might be valuable for better behaved data sets.

Besides speed improvements, our VDAM implementation also replaces inactive classes, which typically leads to higher numbers of suitable classes that better capture the heterogeneity in the data. Compared to the standard SGD algorithm, the VDAM algorithm also shows improved convergence behaviour for 3D initial model generation. Because the VDAM algorithm automatically determines the regularisation parameters, 3D initial model calculations with the VDAM algorithm no longer need to be explicitly limited in resolution, as was the case with the SGD algorithm. Thereby, higher resolution initial models may be calculated without user intervention, which leads to more straightforward selection of suitable initial models. The freedom to progress to higher resolutions may also contribute to better convergence of the VDAM algorithm compared to standard SGD. Although not illustrated here, the VDAM algorithm can also be used for 3D classification and 3D auto-refinement. The latter applications may be particularly interesting in the context of injecting more prior knowledge about protein structures into the 3D reconstruction process [54], which will be a direction of future research.

In previous release of RELION, manual selection of suitable 2D class average images represented a hurdle for automated on-the-fly image processing in the typical workflow. The `relion_class_ranker` program overcomes this hurdle. We found that a combination of a feature vector with a convolutional neural network that acts on the 2D class average images yields excellent results in predicting scores for 2D classes that allow their unsupervised selection. The feature vector contains two types of features. On the one hand, features like the estimated angular and translational alignment accuracy and the class sizes contribute information from the RELION refinement process that is not present in the class average images. On the other hand, hand-crafted features that are calculated from the class average images, such as moments of pixel values in protein and solvent masks, allow biasing the scores on information that is assumed to be important.

The class scores from the `relion_class_ranker` program are on an absolute scale. Although a default selection threshold of 0.15 allowed automated structure determination for eleven out of twelve test data sets, in practice many users may choose to tune the threshold value for their specific type of data. Tuned thresholds gave particle selections with FPRs and FNRs of less than 0.25 for all data sets tested. Ordering 2D class averages by their predicted class scores may also be useful for manual selection. In previous releases of RELION, 2D class average images were typically displayed sorted on their class size. However, the VDAM algorithm often converges to solutions that also contain relatively large classes with suboptimal particles. Therefore, we have observed that sorting the classes based on their predicted scores is also helpful for manual selection of suitable 2D classes. Executing `relion_class_ranker` typically takes less than a minute.

The development of Schemes for the automated execution of pre-defined, image processing workflows that represent branched decision trees is a less visible part of the improvements in RELION-4.0. As an example of what is possible, we distribute the `prep` and `proc` Schemes for automated structure determination inside the new `relion_it.py` approach. Although we show the usefulness of this fully automated approach on twelve test data sets, we expect that many users will want to modify parts of this approach to fit their specific needs. The Schemes are aimed at providing the flexibility that will be required by the different types of end-users to automate a wide range of image processing tasks.

In general, as cryo-EM structure determination continues to improve rapidly, we envision that flexibility in the design of image processing workflows will remain essential for many users. The RELION tools described here aim to facilitate this flexibility, as well as speed, and to help the inexperienced user in getting the most of their cryo-EM images, while at the same time providing the advanced user with all the tools necessary to solve the most difficult structures. Moreover, by distributing these tools as free, open-source software, we encourage the cryo-EM community to build on the advances described.

## Acknowledgements

We are grateful to Matthew Iadanza, Colin Palmer and Tom Burnley for helpful discussions, and to Jake Grimmett and Toby Darling for help with high-performance computing. This work was funded by the UK Medical Research Council (MC_UP_A025_1013 to SHWS) and the European Union’s Horizon 2020 research and innovation programme (under grant agreement 895412 to DK).

## Data availability

RELION-4.0 is distributed under a GPLv2 license and can be downloaded for free from https://github.com/3dem/relion/tree/ver4.0. The 2D class average images used for training the neural network in the `relion_class_ranker`, together with all necessary metadata to replicate the training, can be downloaded from the EMPIAR data base (entry-ID 10812).

## Notes

https://github.com/3dem/relion/tree/ver4.0

https://relion.readthedocs.io/en/release-4.0

### Competing Interest Statement

The authors have declared no competing interest.

